# Simple and rapid separation of diverse neoagaro-oligosaccharides

**DOI:** 10.1101/363739

**Authors:** Fudi Lin, Yayan Huang, Na Zhang, Jing Ye, Meitian Xiao

## Abstract

A rapid and simple method for obtaining pure and well-defined oligosaccharides was established by hydrolyzing agar with β-agarase from *Vibrio natriegens*. The conditions for enzymolysis were optimized as follows: temperature of 45 °C, pH of 8.5, substrate concentration of 0.3%, enzyme amount of 100 U/g and enzymolysis time of 20 h. Neoagaro-oligosaccharides with different degree of polymerizations were gained by hydrolyzing agar with β-agarase at different enzymolysis time. After removing pigment by activated carbon and salts by dialyzing, the enzyme hydrolysis solution was separated with Bio-Gel P2 column chromatography. Neoagaro-oligosaccharides with different degree of polymerizations were acquired. By comparing with standard substances, along with further confirmation by FTIR, MS and NMR, structures of the purified neoagaro-oligosaccharides were identified as neoagarobiose, neoagaroteraose, neoagarohexaose, neoagarooctaose, neoagarodecaose and neoagarododecaose with purities more than 97.0%, respectively. The present study established a method for rapid preparation of various monomers of neoagaro-oligosaccharides that may be of great significance for further study of bioactivities.

## Introduction

Agar, an important marine polysaccharide extracted from the cell walls of red algae, is a linear polymer containing (1→4)-linked 3,6-anhydro-α-L-galactose and (1→3)-linked β-D-galactopyranose [1], and it is composed of agarose and agaropectin [2]. Agar is widely used in food, biological, and pharmaceutical industries [3]. Accumulating reports have indicated the oligosaccharides prepared from agar/agarose have diverse physiological functions such as antioxidant [4-8], anti-hyperlipidemia [9, 10], anti-inflammation [11-13], and whitening effect [14, 15] etc, which will expand its use in the food, cosmetic, and medical industries.

Generally, oligosaccharides from agar/agarose can be classified into agaro-oligosaccharides (AOS) [16-18] and neoagaro-oligosaccharides (NAOS) [19-21], the former are the hydrolysis products of acid or α-agarase which cleaves α-(1→3)-galactosidic bond of the polysaccharides, and the latter are hydrolysates of β-agarase which splits β-(1→4) bond. AOS and NAOS with different degree of polymerizations (DPs) have been reported to possess various bioactivities. A large amount of reports indicated that AOS have the beneficial effects of antioxidant [4-8], anti-obesity [9, 10], anti-inflammation [11-13], anti-cancer [22], and protect the intestine [9, 10, 22] etc. In recent years, NAOS mixtures as well as NOAS of signal DP have obtained increasing attention for its distinct physiological and biological activities. According to the reported prebiotic studies, the NAOS with DPs of 4-12 could increase the amount of *lactobacilli* and *bifidobacteri in vivo*, suggesting they had good probiotic effect [23]. Neoagarobiose (NA2), neoagaroteraose (NA4), and neoagarohexaose (NA6) were reported to have *in vitro* skin whitening and moisturizing effects, among them, NA4 was found to be a better whitening agent than the other two, whereas AOS did not exhibit the same activities[14, 15]. Moreover, NA4 displayed better ability to scavenge hydroxyl radicals compared with NA2, NA6, and neoagarooctaose (NA8), meanwhile, the scavenging activities on hydroxyl radicals was not found for agarotriose and agarobiose. Additionally, NA4 was proved to inhibit inflammation in LPS-stimulated macrophages through suppression of MAPK and NF-κB pathways [24, 25].

Notably, these studies mainly focused on the activities of NAOS mixture, NA2, NA4, and NA6, while few reports investigated the bioactivities of NAOS with higher DPs such as NA8, neoagarodecaose (NA10), and neoagarododecaose (NA12), etc. The reasons may be attributed to the difficulty of obtaining NAOS with higher DPs and the complexity of purification of them. Many efforts have been devoted to obtain NAOS. β-Agarases from marine bacterium *Janthinobacterium* sp. SY12, *Vibrio* sp. Strain JT0107, *Agarivorans albus* YKW-34 and *Agarivorans albus* OAY02 were employed to degrade agarose to give NA2 and NA4 [26-28]. And β-agarase obtained from *Microbulbifer* sp. Q7, marine *Alteromonas* sp. SY37–12, *Pseudoalteromonas* sp. CY24, marine *Agarivorans* sp. LQ48 and *Pseudoalteromonas* sp. AG4 could hydrolyze β-1,4-glycosidic linkages of agarose/agar to gain NA4 and NA6 [29-33]. β-Agarases isolated from *Stenotrophomonas* sp. NTa and *Agarivorans* sp. JA-1 in *Bacillus subtilis* degraded agarose/agar to NA2, NA4 and NA6 as the predominant products [34, 35]. *Agarivorans albus* OAY02 could secrete two β-agarases, among them, one β-agarase could cleave agarose into NA2 and NA4, the other β-agarase made agarose become NA2, NA4, NA6 and NA8 [36]. And β-agarase from marine *Pseudoalteromonas* sp. CY24 could degrade agarose to NA8 and NA10 [37]. β-Agarase from marine bacteria can degrade agarose to NAOS with different DPs, however, up to now it has not been reported that β-agarase could degrade agarose/agar to obtain NAOS with desired DPs at different enzymolysis time.

Developing methods for rapid separation and purification of NAOS is also very important for obtaining purified NAOS. Size-sieving based on gel-permeation chromatography (GPC) and high performance anion exchange chromatography (HPAEC) are commonly used methods for the separation of polysaccharides and oligosaccharides [6, 38-41]. Toyopearl HW-40S was used to purify NA4[38]. It was reported that NAOS with DP of 2, 4, and 6 could be separated and purified by Bio-Gel P2 [40, 41] and NA4, NA6, NA8, NA10 and NA12 could be separated by two chromatography steps of consecutive Bio-Gel P-6 chromatography [40]. Sephadex G-10 combined with G-25 was employed to purify AOS [6, 39]. The SEC-HPLC and NH_2_-HPLC systems were used to isolate and purify NAOS and AOS [42]. A HPAEC system equipped with a semi-preparative CarboPac™ PA100 column was applied to prepare NAOS and AOS from DP 2 to DP 22 with product yield and purity no more than 17.2% and 77.7%, respectively [43]. However, these methods are a little bit complicated, time-consuming, and instrument-depending, and the product yield and purity of obtained oligosaccharides are not high enough for further study. Therefore, to deeply understand the bioactivities and the mechanism of NOAS, it is urgent to develop a simple and rapid method for the preparation of NAOS with various DPs.

In the present study, a simple method of obtaining NAOS with desired DPs was established by regulating the enzymolysis time of β-agarases. And a gel filtration chromatography was developed for the rapid preparation of each NAOS with different DPs in high quality and quantity, making it possible to the further studies on their bioactivities.

## Materials and methods

### Strains and reagents

NA2, NA4, NA6, NA8, NA10 and NA12 used as standards were purchased from Qingdao bozhihui Biological Technology Co. Ltd (Qingdao, China). Acetonitrile were gained from Sigma-Aldrich (St. Louis, MO, USA). All the other reagents were commercially available and of analytical grade.

The culture and fermentation condition of *Vibrio natriegens* was the same as what was explored in our previous work[44]. The obtained β-agarase was purified from the fermentation liquor with the combination of ammonium sulfate salting-out, dialysis, ion exchange chromatography and gel-filtration. The enzymatic activity of the purified β-agarase reached 103 U/mL, which was applied in this work for the later experiment.

### Optimization of enzymatic hydrolysis condition

The enzymatic hydrolysis condition including reaction temperature (30, 35, 40, 45, 50, 55 and 60 °C), pH (6.0, 6.5, 7.0, 7.5, 8.0, 8.5 and 9.0), reaction time (1, 5, 10, 15, 20, 25 and 30 h), substrate concentration (0.1, 0.2, 0.3, 0.4, 0.5, 0.6 and 0.7%) and enzyme amounts (20, 40, 60, 80, 100, 120 and 140 U/g) were optimized. All hydrolysis reactions were conducted in triplicate.

### Preparation of NAOS products with different enzymolysis time

3% Agar solution was completely dissolved in 1 L of 0.1 M Tris-HCl solution by heating and then cooled to 45 °C. Subsequently, the solution was treated with the β-agarase for 4 h, 6 h, 8 h, 10 h and 12 h to obtain products A, B, C, D and E, respectively. After inactivation with boiling water bath for 15 minutes and centrifugation for 30 min (12,000 × g), the insoluble agar was removed and the NAOS in supernatant was filtered with a 0.22 μm membrane (Millipore, Cork, Ireland). The crude products of different enzymolysis time were finally obtained by lyophilization and stored at −20 °C for further use.

### Analysis of NAOS products by HPLC-ELSD

The NAOS products were analyzed by HPLC-ELSD system which consisted of a Waters e2695 HPLC system (Waters, Milford, MA, USA) equipped with an evaporative light scattering detector (Waters 2424, USA). Separation was performed on an Asahipak NH_2_P-50 4E multi-mode analytical column (250 mm×4.6 mm, 5 μm) with the column temperature of 30 °C. Isocratic elution was conducted with acetonitrile-water (65:35) as the mobile phase with a flow rate of 1 mL/min. The injection volume was set at 10 μL and the detector nebulizer temperature was 75 °C.

### Rapid purification of NAOS

For the purification of NAOS, 1 g crude NAOS product powder was resuspended in 200 mL distilled water, followed by addition of 10 g activated carbon. The mixture was stirred for 2 h and then NAOS were washed out by 30% ethanol solution and ethanol was removed by rotary evaporator. Then the left was freeze dried and detected.

GPC was applied to separate the NAOS. 250 mg of products A was resuspended in 1 mL of NH_4_HCO_3_ (0.1 M), and the solutions were loaded onto the Bio-Gel P2 column (1.8 × 150 cm, Bio-Rad Laboratories, Hercules, CA, USA), respectively. NH_4_HCO_3_ was used as eluent at a flow rate of 0.4 mL/min and fractions of 4 mL each were collected. Then the collected fractions were detected by TLC on a Silica Gel 60 plate (Merck, Darm-stadt, Germany) developed with a solvent of isopropanol/water/ammonium hydroxide (30:15:2, v/v/v). After being sprayed by anisaldehyde and heated for 10 min, the spot of the products could be visualized.

### Identification of NAOS

The structure of isolated NAOS was elucidated by FTIR and the molecular mass was confirmed by ESI-TOF-MS analysis. The FTIR spectra of KBr pellets of the NAOS after drying at 105 °C for 2 h were recorded on the FTIR (FTIR-84, Shimadzu, Japan) spectrophotometer. Scans were performed over the range of 4000 - 400 cm^−1^ with a resolution of 4 cm^−1^ for 32 times. ESI-TOF-MS analysis was carried out on a Q Exactive Hybrid Quadrupole Orbitrap mass spectrometer (Thermo, Bremen, Germany) coupled with an ESI source in positive ion mode. Recorded mass range was *m/z* 200 - 2000. 1H-NMR and 13C-NMR spectra were measured in D_2_O on a Avance 500 spectrometer (Bruker, Avance III, Switzerland, 500 MHz 1H, 125 MHz 13C) at room temperature and C_3_D_6_O was added as an internal standard with the chemical shifts were reported in δ = 2.05 ppm for 1H-NMR, δ = 29.84 ppm for 13C-NMR.

## Results and discussion

### Optimization of hydrolysis parameters by single factor experiments

Single factor experiments were carried out to optimize the enzymatic hydrolysis process. Fig. 1a displayed the influence of temperature on the yield of reducing sugar. Temperature is an important factor in the process of enzymatic hydrolysis. Within a certain temperature range, the raise of temperature is beneficial to enzymatic reaction. As shown in Fig. 1a, the yield of reducing sugar was increased with the elevation of temperature and it reached 45.2% at the temperature of 45 °C, after that the yield was decreased significantly with the raise of temperature. Therefore, the temperature was set at 45 °C. The pH is another vital parameter in the enzymatic hydrolysis process, because enzyme activity is greatly affected by pH value. Generally, an enzyme can maintain high enzyme activity under suitable pH. As known in Fig. 1b, the yield of reducing sugar was increased with the increase of pH from 6.0 to 8.5, the largest yield of reducing sugar was 42.7% at pH 8.5. Then the yield was decreased, which may be attributed to the increase of pH on the inhibition of the β-agarase activity. As for the substrate concentration, low substrate concentration leads to low enzyme utilization, however, too high the substrate concentration will hinder the diffusion of molecules and reduce the enzymatic reaction rate. As showed in Fig. 1c, when the substrate concentration was set at 0.3%, the yield of reducing sugar reached the highest of 43.3%. In addition, not only enzyme amount but also enzymolysis time has prominent effects on the yield of reducing sugar [45, 46]. The enzyme amounts were also investigated in the present study and the adding amount at 100 U/g turned out to be optimal with the reducing sugar yield of 42.4%, and the yield of reducing sugar had remained almost stable after the enzyme amount added was more than 100 U/g. Fig. 1e indicated that the yield of reducing sugar increased with the extension of enzymolysis time before the time of 20 h. When the enzymolysis time was 20 h, reducing sugar yield reached the highest at 43.3%, and the generation of reducing sugar didn’t appear to increase over 20 h of enzymolysis time. Therefore, the optimized conditions were optimized as follows: the temperature of 45 °C, the pH of 8.5, the substrate concentration of 0.3%, the enzyme amount of 100 U/g and the enzymolysis time of 20 h.

**Fig. 1.**
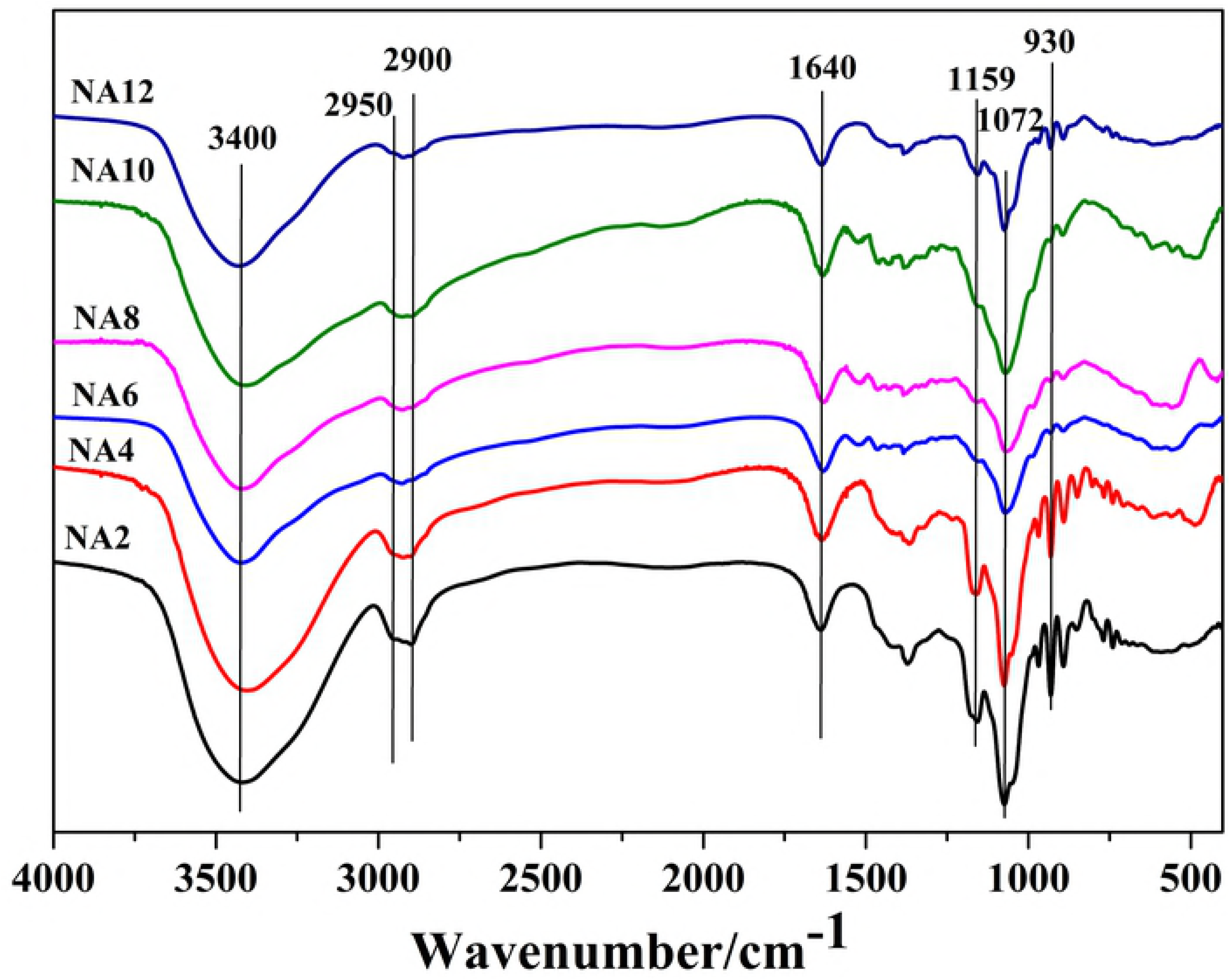
Effects of enzymolysis condition on reducing sugar yield

### Preparation of NAOS with different DPs

The β-agarase obtained by the different strains would degrade agar to gain the NAOS with different DPs [26, 30, 34, 35]. Specifically, the regular changes of NAOS with different DPs at different enzymolysis time were found in the present study (Fig.2), which was not mentioned in the previous reports. The enzymolysis products were determined with HPLC-ELSD by comparing with standard substances and the content of each oligosaccharide was calculated from the regression equations, as shown in Fig.2 and table 1 and table 2, and the enzymolysis product was analyzed to be composed of NA2, NA4, NA6, NA8, NA10 and NA12 with the yield of 5.0%, 38.9%, 18.1%, 16.8%, 13.8% and 2.5% respectively after the agar was hydrolyzed by β-agarase for 4 h. When the hydrolysis time was 6 h, NA12 was completely hydrolyzed and the product obtained was NA2, NA4, NA6, NA8, and NA10, the yield were 6.0%, 52.7%, 15.2%, 13.7%, and 11%, respectively. As the hydrolysis time prolonged to 8 h, NA10 was not detected in the product and the yield for NA2, NA4, NA6 and NA8 were 9.3%, 59.5%, 13.7% and 11.7%, respectively. With the extension of the hydrolysis time, the DPs of NAOS obtained were getting smaller and smaller. When the hydrolysis time was 10 h, the product was consisted of NA2, NA4, and NA6 with the yield of 13.8%, 68.4% and 9.6% separately. And the hydrolysis time reached 12 h, only NA4 and NA2 were left in the product with the yield of 21.6% and 71.1%, and the product components were no longer changed with the extension of time.

**Fig. 2.**
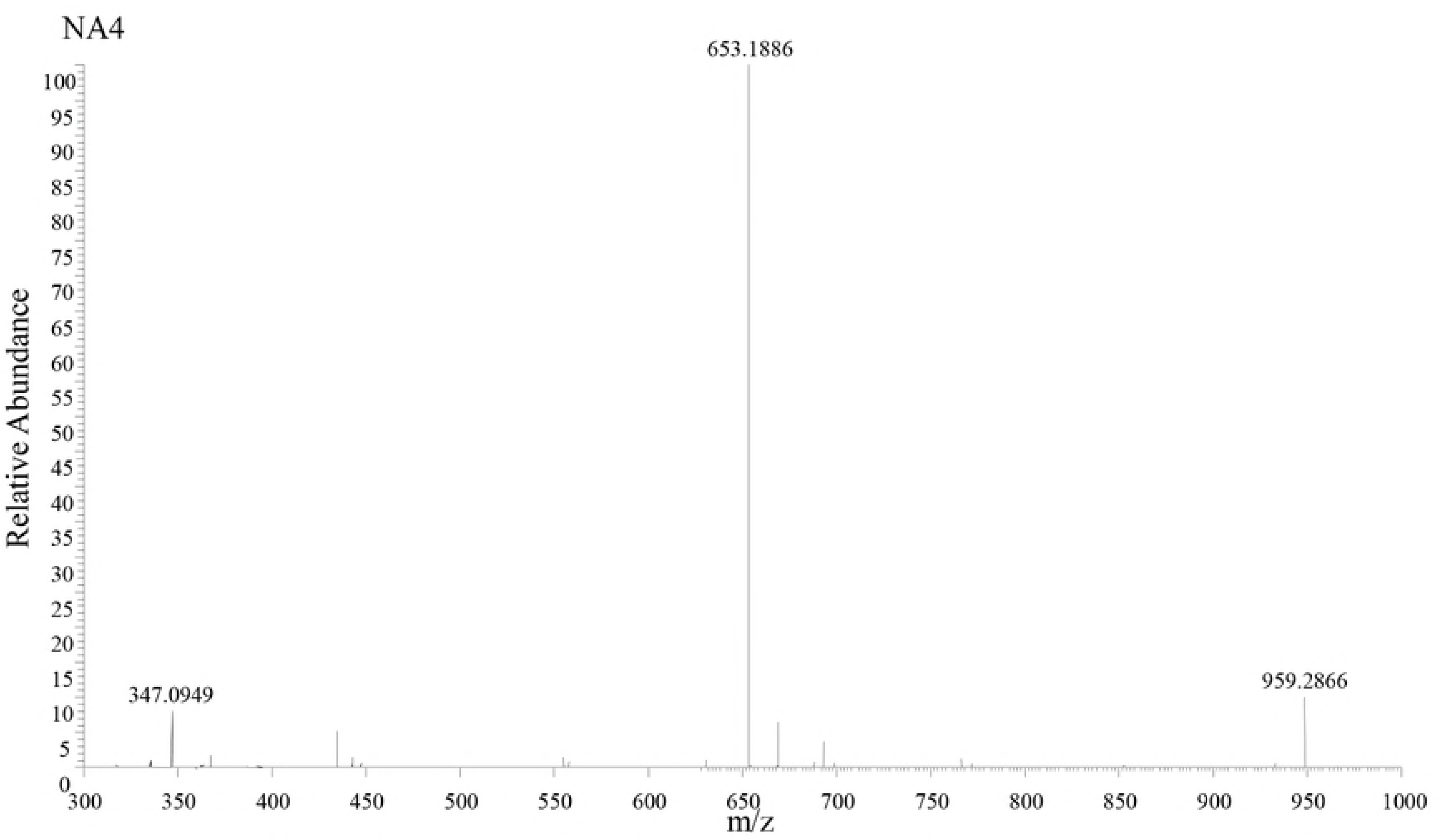
Anlysis of enzymatic hydrolysates with different enzymolysis time, S-NAOS standards, 1-NA2, 2-NA4, 3-NA6, 4-NA8, 5-NA10, 6-NA12; A 4 h, B 6 h, C 8 h, D 10 h, E 12 h

**Table. 1.** The regression equations of NAOS.

| Oligosaccharide | Equation | R<sup>2</sup> |
| --- | --- | --- |
| NA2 | y=6.29305x-2.21022 | 0.992 |
| NA4 | y=6.05888x-3.18131 | 0.996 |
| NA6 | y=5.54881x-3.71928 | 0.997 |
| NA8 | y=4.8549x-4.42057 | 0.996 |
| NA10 | y=4.48493x-3.88547 | 0.994 |
| NA12 | y=4.25971x-3.75222 | 0.992 |

x: the concentration of NAOS (mg/ml), y: peak area $\times 10^{-6}$ .

**Table. 2.** The percentage of monomers at different hydrolysis time.

|  | A | B | C | D | E |
| --- | --- | --- | --- | --- | --- |
| NA2 (%) | 5 | 6 | 9.3 | 13.8 | 21.6 |
| NA4 (%) | 38.9 | 52.7 | 59.5 | 68.4 | 71.1 |
| NA6 (%) | 18.1 | 15.2 | 13.7 | 9.6 | 0 |
| NA8 (%) | 16.8 | 13.7 | 11.7 | 0 | 0 |
| NA10 (%) | 13.8 | 11 | 0 | 0 | 0 |
| NA12 (%) | 2.5 | 0 | 0 | 0 | 0 |

At present, there are mounting reports on the preparation of NAOS by enzymatic method and agarase is mainly derived from the secretion of marine bacteria. Frequently β-agarase is found to degrade agar to generate NA2, NA4, or NA6 as the main products. The DagA secreted β-agarase which could hydrolyze agar to gain NA2, NA4 and NA6 [47]. And it was reported β-agarase from *Stenotrophomonas sp.* NTa degraded agar only obtained NA2, NA4 and NA6 as the predominant products and a small amount of 3,6-anhydro-α-L-galactose, and notably the products did not change with the change of the hydrolysis time [35]. At the same time, the composition of the agarolytic did not changed over time by some β-agarase, which included four even-numbered NAOS with DP of 2-8, and the amount of NA4 was more than others [25]. However, these methods found in literature were used in preparing NAOs mainly with DP no more than eight [25, 35, 47]. Interestingly, we found in our study that the DPs of the NAOS reduced regularly with the enzymolysis time increased every two hours, and the final product was composed of NA4 and NA2. Therefore, desired NOAS with different DPs could be obtained by controlling the hydrolysis time, which was beneficial to the further studies of NAOS.

### Rapid isolation of NAOS

Separation and purification of NAOS was carried out using a Bio-Gel P2 column and detected by TLC, and the result was shown in Fig. 3. For product A, fractions of 18 to 22 were NA12 with the yield of 3.2%, fraction 24 to fraction 27 were NA10 with the yield of 4.2%, fraction 30 to fraction 35 was NA8 with the yield of 7.5%, fraction 37 to fraction 40 was NA6 with the yield of 10.2%, fraction 44 to fraction 56 was NA4 with the yield of 35.8%, fraction 61 to fraction 64 was NA2 and the yield was 23.2%, respectively. After detected by HPLC-ELSD, the purity of NA2, NA4, NA6, NA8, NA10 and NA12 were 99.3%, 98.9%, 98.0%, 97.6%, 97.3% and 97.4%, respectively (Fig.4).

**Fig. 3.**
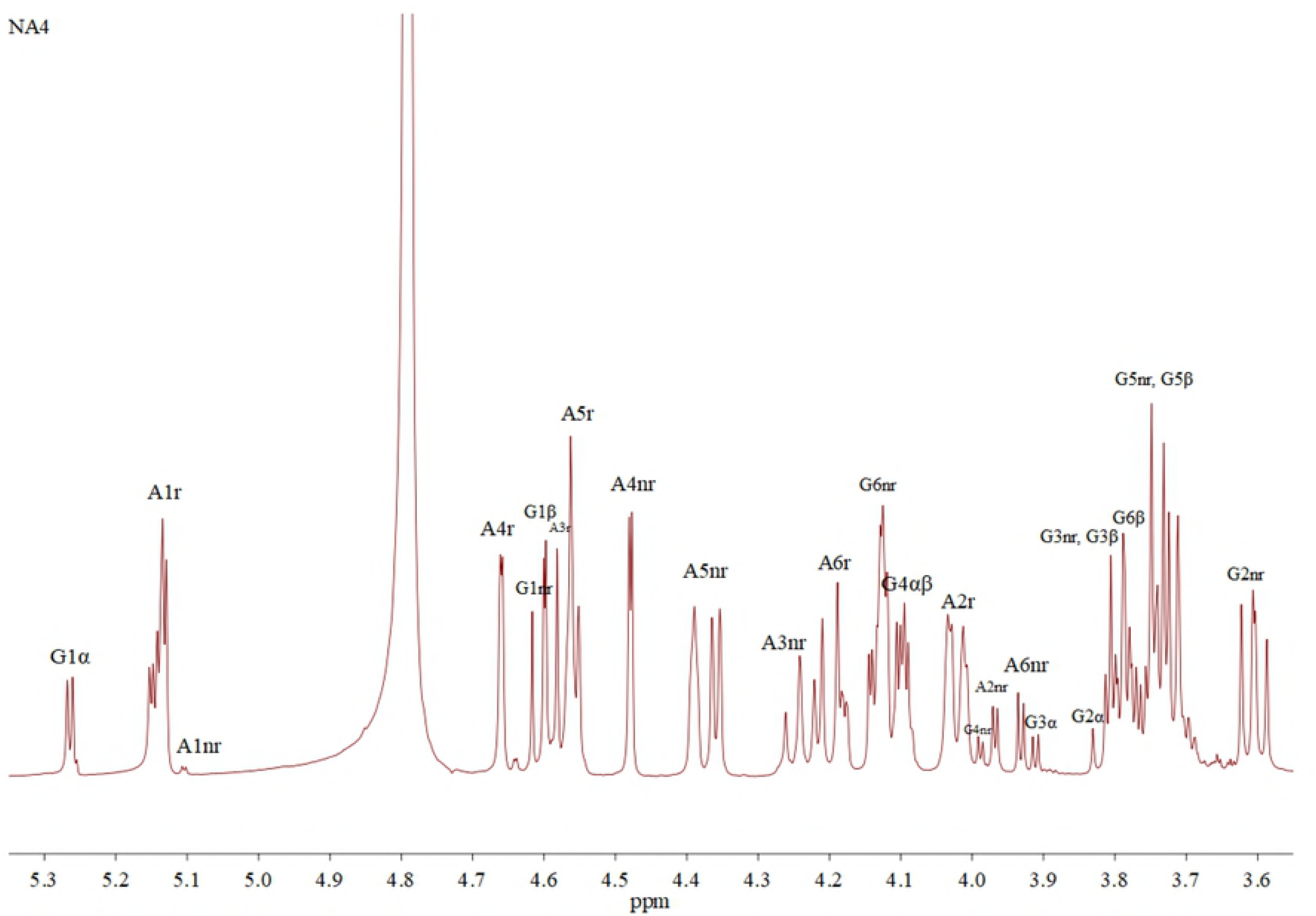
TLC analysis of purified NAOS. The ladder of NAOS with different DP (2-12); S: NAOS standards; Fractions 18-22: NA 12, Fractions 24-27: NA 10, Fractions 30-35: NA8, Fractions 37-40: NA 6, Fractions 44-56: NA4, Fractions 61-64: NA2

**Fig. 4.**
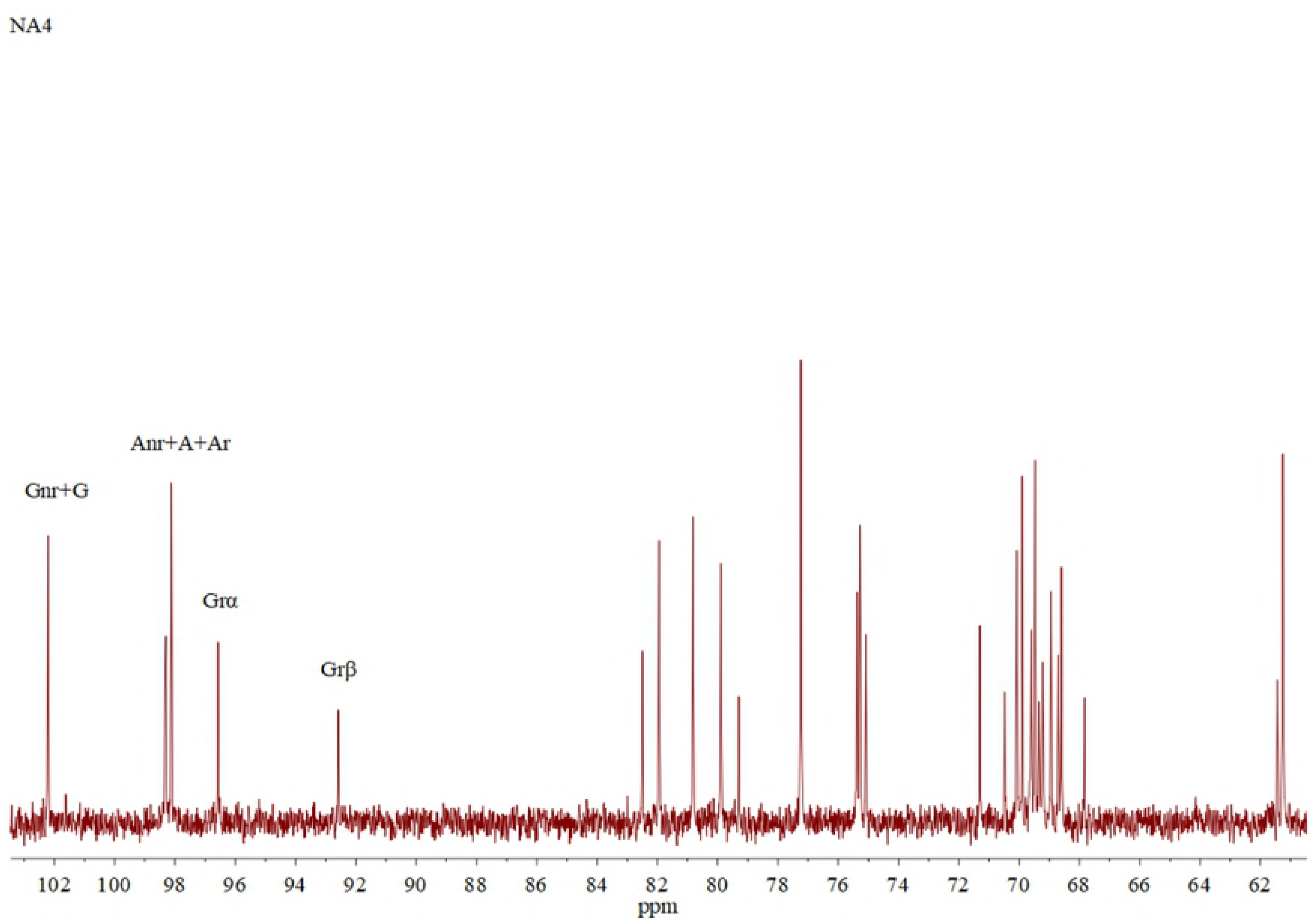
High liquid chromatograms of purified products

GPC and HPAEC are commonly used method for separation and preparation of NAOS and Bio-Gel P-2 and Bio-Gel P-6 are frequently applied to purify NAOS [6, 38-43]. However, these two separation media are usually jointly used to prepare NAOS with diverse DPs. Noticeably, in the present study, one chromatography step of Bio-Gel P-2 column chromatography could be developed to obtain NAOS of DP2-12 with purities more than 97%, suggesting it is a simple and rapid method for the preparation of NAOS.

### Structure and molecular confirmation of NAOS

The structures and the molecular mass of the purified NAOS were confirmed by FTIR and MS analysis. Fig.5 demonstrated the results of the FTIR analyses. In all the six obtained oligosaccharides, the disappearance of absorption band around 1260 cm^−1^ indicated the elimination of sulfate group in the degradation process. There was a broad absorption band around 3400 cm^−1^, which may be assigned to hydroxyl group. The region around 2950 cm^−1^ and 2900 cm^−1^ were assigned to C-H. The band around 1640 cm^−1^ suggested the existence of C-C sugar ring. The fingerprint region, including many FTIR absorptions of specific characteristic bonds, was a region of lower wave numbers. There was absorption band around 1159 cm^−1^ was the stretch vibration of C-O within C-O-H. Absorption bands appeared at 1072 cm^−1^, indicating the presence of C-O within C-O-C bond. A well-defined peak was shown at about 930 cm^−1^ corresponding to 3, 6-anydro-D-galactose.

**Fig. 5.**
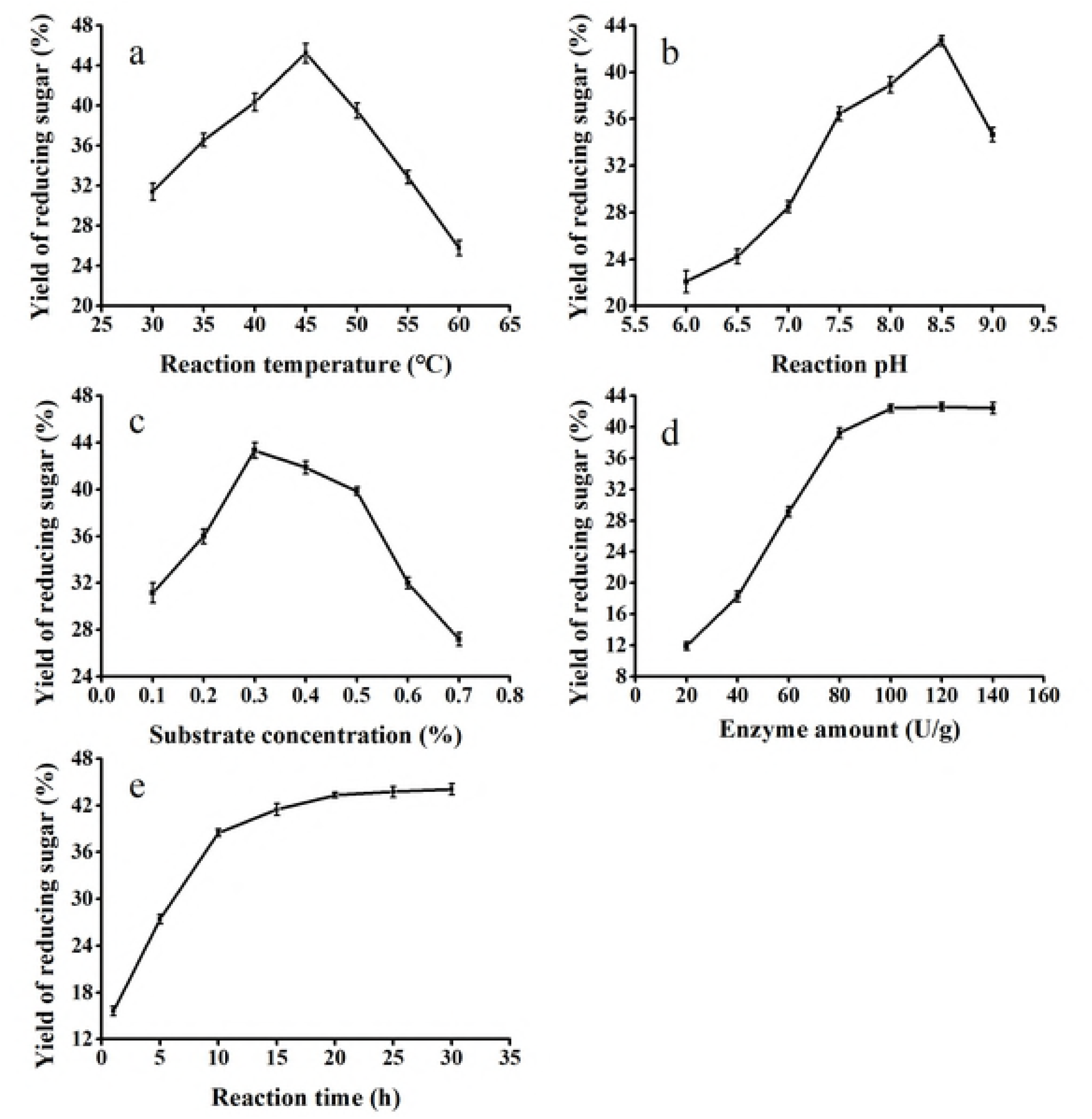
FTIR spectrum of NAOS

The ESI-TOF-MS analysis results were displayed in on table 3, confirming the purified oligosaccharides were NA2, NA4, NA6, NA8, NA10 and NA12, respectively. The NA4 was shown in Fig. 6 and other monomers were in S1 Fig.

**Fig. 6.**
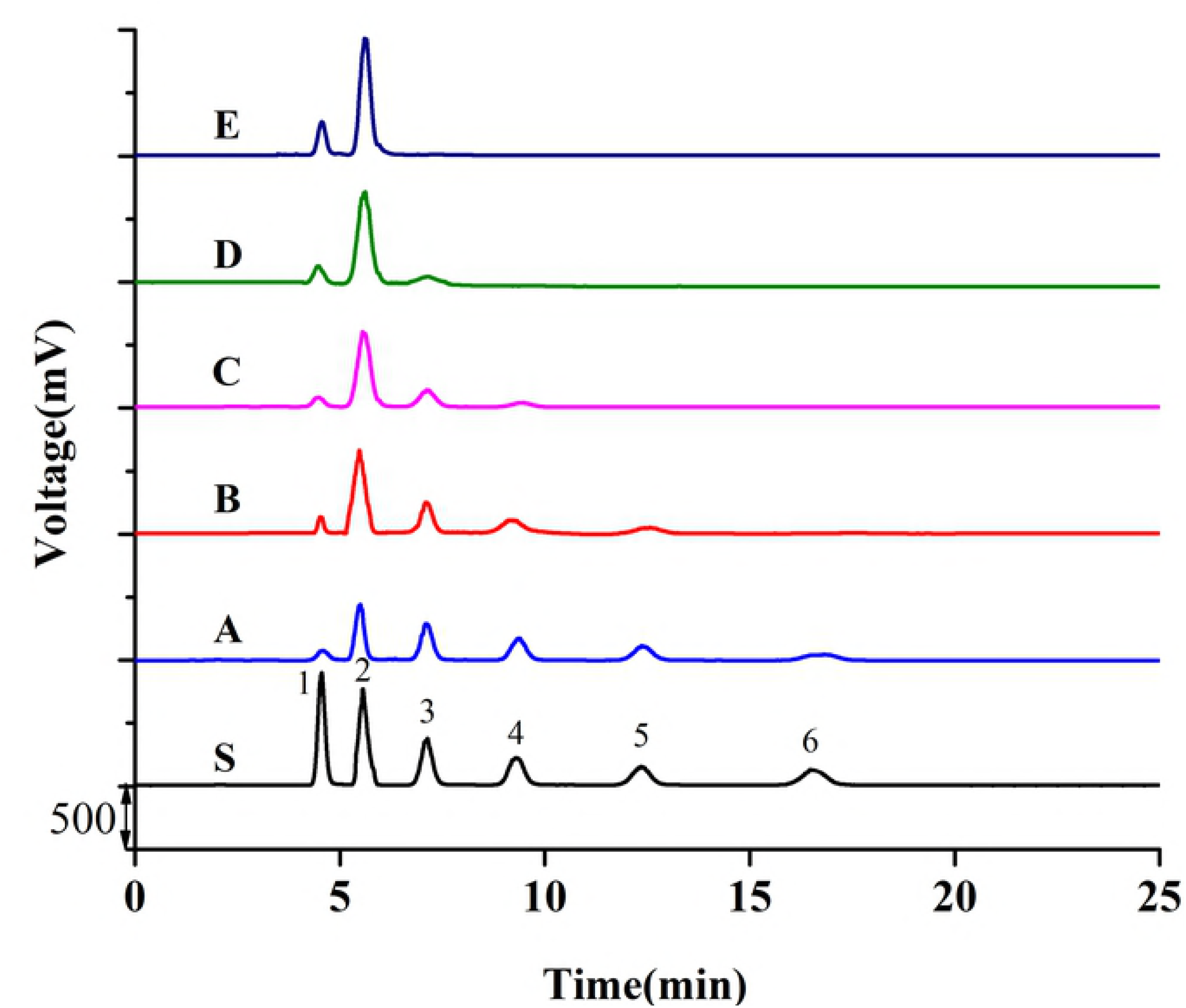
The ESIMS spectra of NA4

**Table 3.** The ESI-TOF-MS analysis of the purified products.

| purified products | MS signals(m/z)<br>[M+Na]<sup>+</sup> | Calculated molecular<br>weights |
| --- | --- | --- |
| NA2 | 347.0949 | 347.0954 |
| NA4 | 653.1886 | 653.1905 |
| NA6 | 959.2866 | 959.2856 |
| NA8 | 1265.3752 | 1265.3807 |
| NA10 | 1571.4752 | 1571.4757 |
| NA12 | 1877.5751 | 1877.5708 |

The structural information of six monomers was demonstrated by ^1^H-NMR and ^13^C-NMR spectroscopy. Assignments of ^1^H-NMR and^13^C-NMR spectroscopy were built the close similarity with literature values, and the interpretation of these signals was shown in Fig. 7 and table 4 [1, 41, 42]. It indicated twelve particular major anomeric carbon signals (G and A), which was anticipated that the NA2 was the major repeat unit, and the signals (Gnr and Anr) were the residues towards the nonreducing end of the NAOS. Resonances at about 96.6 and 92.6 ppm were characteristic of β and α anomeric form of galactose residues at the reducing end of the NAOS[40, 48], respectively, and the NA4 of ^13^C-NMR was showed on Fig. 8, and ^1^H-NMR and ^13^C-NMR of other monomers were in S2 Fig and S3 Fig, respectively.

**Fig. 7.**
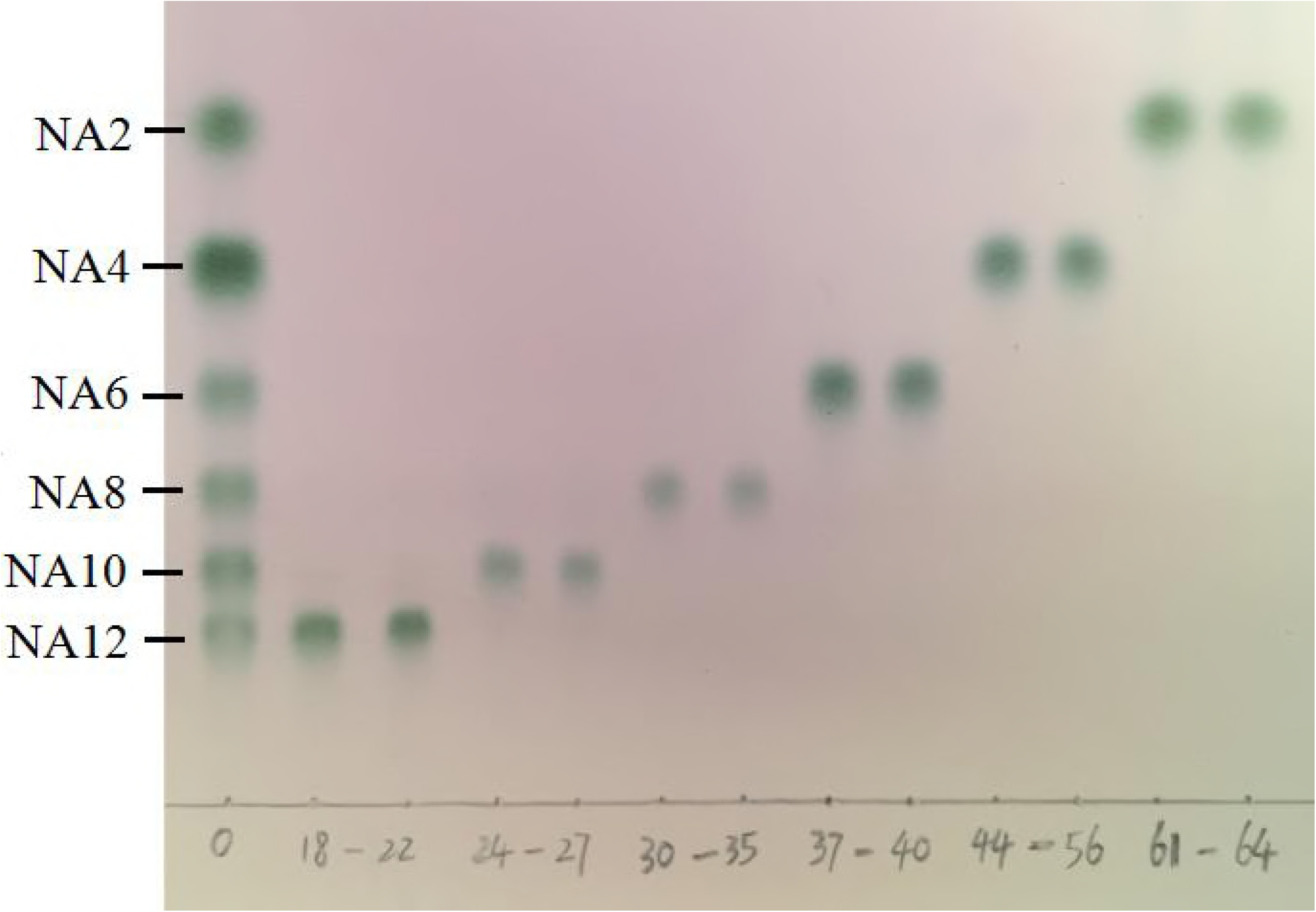
^1^H NMR spectra of NA4, Peak labels: A, 3,6 anhydrogalactose; G, galactose; nr and r refer to the non-reducing and reducing ends; α, β refer to positions of protons on reducing ends; numbers from 1 to 6 refer to place of protons

**Fig. 8.**
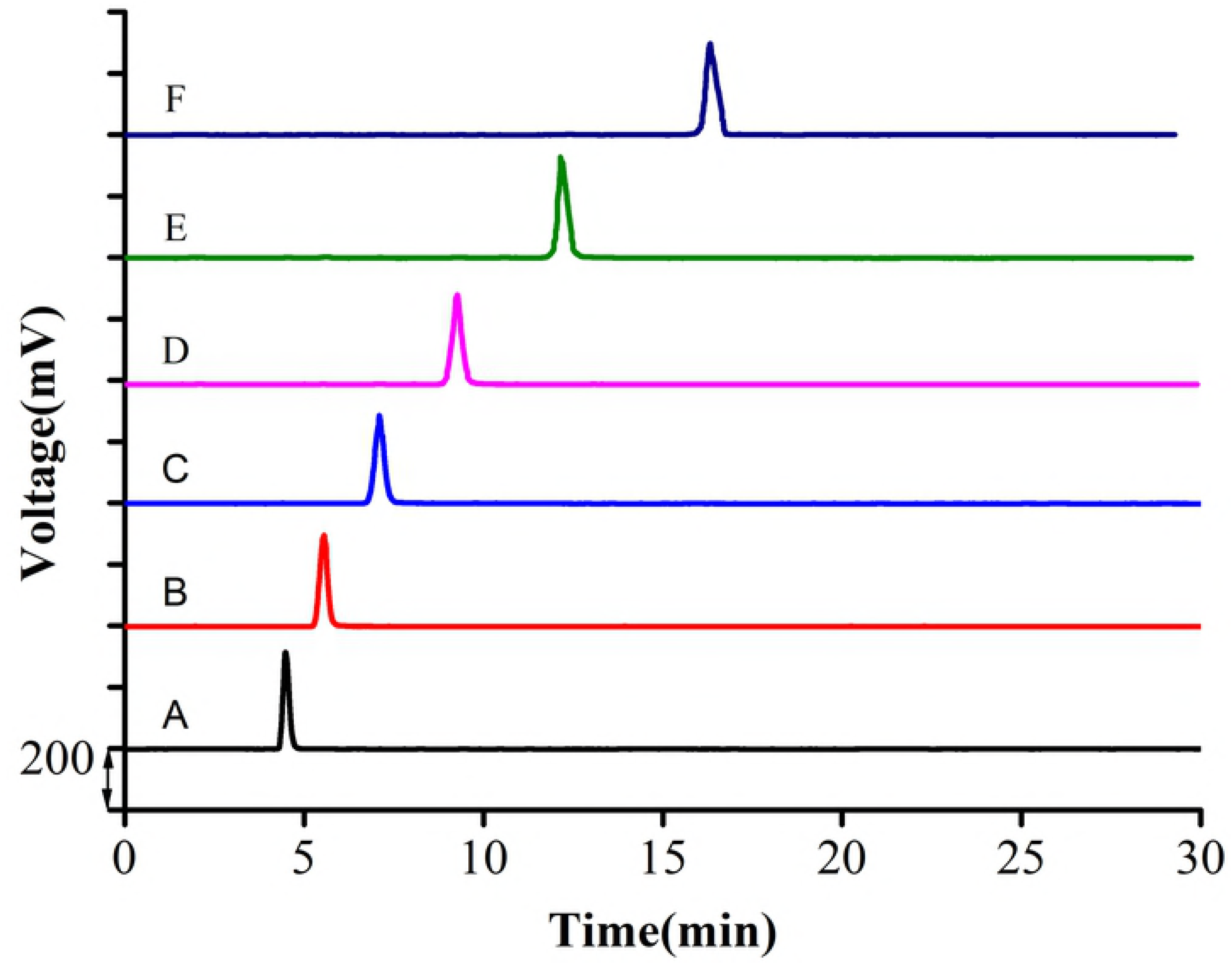
^13^C NMR spectra of NA4

**Table 4.** Chemical shift assignments for ^13^C-NMR spectra of NAOS.

| Unit | C1 | C2 | C3 | C4 | C5 | C6 |
| --- | --- | --- | --- | --- | --- | --- |
| Gnr | 102.2 | 69.9 | 82.0 | 67.8 | 75.1 | 61.3 |
| Grβ | 96.6 | 69.9 | 82.5 | 68.6 | 75.1 | 61.3 |
| Grα | 92.6 | 67.8 | 79.9 | 69.2 | 69.9 | 61.4 |
| Anr | 98.3 | 69.4 | 80.8 | 69.6 | 77.3 | 68.7 |
| Arα | 98.1 | 69.5 | 79.9 | 77.3 | 75.3 | 69.0 |

## Conclusions

In summary, the present study has developed a feasible approaches for the preparation of desired NAOS with different DPs by regulating the enzymolysis time of β-agarase. Furthermore, the NAOS of diverse DPs were rapidly and simply prepared by the Bio-gel P2 column chromatography with purities higher than 97% for further evaluating their bioactivity potentials.

## Acknowledgment

This work was supported by Subsidized Project for Postgraduates’ Innovative Fund in Scientific Research of Huaqiao University; and the Public Science and Technology Research Funs Projects of Ocean [grant numbers 20130515-2, 201505026-5].

## Supporting information

**S1 Fig. The ESIMS spectra of NA2, NA6, NA8, NA10 and NA12**

**S2 Fig. ^1^H NMR spectra of NA2, NA6, NA8, NA10 and NA12**

**S3 Fig. ^13^C-NMR spectra of NA2, NA6, NA8, NA10 and NA12**

